# TB-Bench: A Systematic Benchmark of Machine Learning and Deep Learning Methods for Second-Line TB Drug Resistance Prediction

**DOI:** 10.64898/2026.04.08.717138

**Authors:** Brintha VP, Saish Jaiswal, Ansh Meshram, Deepti PVS, Sidharthan S C, Manikandan Narayanan

## Abstract

Drug-resistant tuberculosis (TB), characterized by prolonged treatment regimens and suboptimal treatment outcomes, remains a major obstacle to global TB elimination. Advances in sequencing technologies have enabled the development of machine-learning (ML) approaches, including deep-learning (DL) methods, to predict drug resistance directly from genomic data. However, a significant gap remains in translating these advances into clinical practice. While current approaches reliably predict resistance to first-line drugs, they show consistently lower and more variable performance for second-line drugs compared with traditional drug-susceptibility testing. To characterize these limitations and assess practical utility, we conducted a comprehensive survey and standardized benchmarking of current approaches for predicting TB drug resistance using whole-genome sequencing (WGS) data. Using systematic selection criteria, we identified 20 traditional ML and DL models from 8 studies and evaluated drug-specific versions across 14 second-line drugs within a unified framework. To account for methodological heterogeneity, the models were evaluated using three distinct feature sets reflecting variability in input representations. We trained and evaluated the models on different subsets of the WHO dataset, comprising 50,801 samples, and assessed generalizability using an external validation dataset comprising 1,199 samples. In the internal evaluation on the held-out WHO test dataset, traditional ML models using binary features achieved higher predictive performance than DL models. For example, XGBoost achieved the highest area under the precision-recall curve (PRAUC) scores (46%–93%) for 10 of the 14 drugs. However, performance varied substantially across drugs. Notably, the superior performance of traditional ML models — even with limited feature sets — highlights their applicability in low-resource settings. When evaluated on the external validation dataset, the performance of traditional ML and DL models was comparable, and neither class of models demonstrated substantial improvement over catalogue-based approaches, underscoring challenges in cross-dataset generalization. Overall, this benchmarking study provides a comprehensive and systematic evaluation of current approaches, establishes a rigorous evaluation framework for future comparisons, and identifies key methodological considerations necessary to advance robust drug resistance prediction in clinical settings. To enhance reproducibility and facilitate the application of TB-Bench to additional datasets and models, we have made the source code publicly available at https://github.com/BIRDSgroup/TB-Bench.

## 1 Introduction

Global efforts to eradicate tuberculosis (TB) are hindered not only by prolonged treatment regimens but also by the escalating challenge of drug resistance. According to the 2023 World Health Organization (WHO) report, 10.8 million people were estimated to have TB, of which nearly 400,000 were estimated to have MDR-TB (resistant to the primary first-line drugs isoniazid [INH] and rifampicin [RIF]) or RIF-resistant TB (RR-TB) [1]. While second-line drugs such as bedaquiline (BDQ) and fluoro-quinolones — including moxifloxacin (MFX) and levofloxacin (LFX) — are effective for MDR/RR-TB, they may fail in cases of extensively drug-resistant (XDR) or pre-extensively drug-resistant (pre-XDR) TB [2]. Pre-XDR TB is defined as MDR-TB with additional resistance to a fluoroquinolone, while XDR-TB involves further resistance to at least one Group A drug, such as BDQ or linezolid (LZD). According to the 2024 WHO report, 28,982 individuals were identified with pre-XDR or XDR-TB among those tested [3]. Although genetic mutations and epistatic interactions in *Mycobacterium tuberculosis* (*M. tb*) are major drivers of resistance, the mismanagement of anti-TB drugs also plays a significant role [4]. Therefore, prospectively identifying the optimal drug combination is crucial for maximizing treatment efficacy and preventing the acquisition of further resistance.

The current reference standard for characterizing resistance is culture-based phenotypic drug-susceptibility testing (DST). However, this method is limited by long turnaround times, contamination risks, and variable sensitivity. Molecular approaches such as Polymerase Chain Reaction (PCR), targeted Next-Generation Sequencing (tNGS), and whole-genome sequencing (WGS) offer faster results and high sensitivity for detecting first-line drug resistance [5]. Consequently, computational methods — including catalogue-based tools (e.g., TBProfiler, Mykrobe), and ML/DL approaches have been developed to infer resistance from WGS data. While catalogue-based approaches enable rapid prediction from known mutations, they fail to capture non-linear or epistatic interactions [6]. Traditional ML methods perform comparably to catalogue-based tools for first-line drugs, but their accuracy for second-line drugs remains mixed [7, 8]. Furthermore, both traditional ML and DL models are often constrained by scarcity of genotyped isolates with matched phenotypic data — particularly for second-line drugs — resulting in overfitting and poor generalization [9].

Although several benchmarking studies have evaluated WGS-based drug-resistance prediction methods, most have focused primarily on catalogue-based approaches [10, 11] or relied on datasets with limited sample sizes (<2,000 samples) [12], underscoring the need for assessment on larger, more diverse cohorts. With the availability of comprehensive resources such as the Bacterial and Viral Bioinformatics Resource Center (BV-BRC) and the WHO global collection, it is now feasible to rigorously benchmark prediction models for second-line drug resistance to guide clinical decision-making. Beyond simply assessing performance metrics, evaluating the interpretability of these models is crucial for understanding their capacity to identify novel resistance-associated mutations.

In this study, we present a comprehensive benchmarking of 20 traditional ML and DL methods, training drug-specific models to predict resistance for 14 second-line drugs. We utilized the WHO dataset comprising 50,801 isolates, which underlies the 2023 TB mutation catalogue. To ensure robust performance estimation and hyperparameter tuning, the WHO data were stratified into distinct training, validation, and held-out test sets. Model performance was evaluated using three distinct feature sets to account for genomic representation variability. Furthermore, we assessed generalizability on an independent external validation dataset of 1,199 isolates. Our results on held-out test sets demonstrate that relatively simple approaches, such as Extreme Gradient Boosting (XGBoost) and Logistic Regression (LR), outperform more complex DL models in predicting second-line drug resistance. The standardized framework established here is readily extensible to additional models and datasets, facilitating the systematic evaluation of future methods.

## 2 Results

### 2.1 Study design — selection criteria and pipeline

Benchmarking was performed on 14 second-line drugs (see Figure 1 for drug names) selected based on the availability of resistant samples, for each drug, in the WHO dataset, comprising 50,801 samples. The final dataset comprised 49,266 samples following quality control and filtering (see details in Section 4.1).

**Fig. 1.**
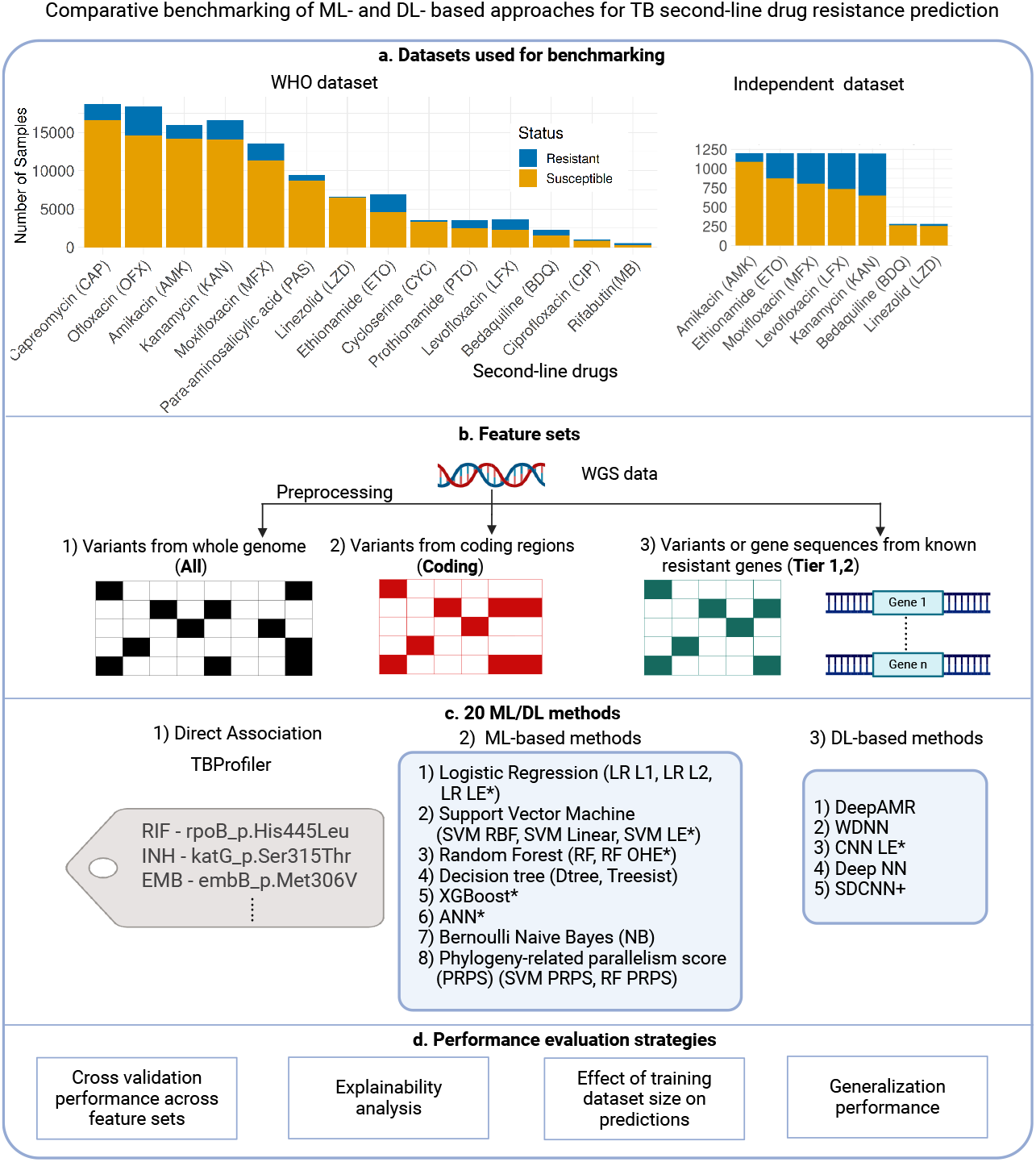
Benchmarking overview: a) The large WHO 2023 TB mutation catalogue dataset was used for model development, and an independent dataset from China was employed to validate the trained models. b) The workflow began with preprocessing WGS data to generate VCF files for each isolate, followed by the construction of input representations for the chosen models. Three distinct feature sets were derived to enable comparative analysis. Here, an asterisk (*) next to a model indicates that the corresponding study employed data balancing through oversampling or downsampling. LE and OHE indicate that the model uses either label-encoded or one-hot-encoded input data, respectively. c) A total of 20 ML/DL methods, chosen from eight published studies, were selected for benchmarking. d) The resulting models were then evaluated on the independent dataset, assessed for performance on unseen drugs, examined for explainability/interpretability, and analyzed to determine the influence of training dataset size on predictive performance.

#### 2.1.1 Method Selection

To ensure a comprehensive evaluation, we conducted a systematic literature search on Google Scholar (as of May 20, 2025) using the query: (“tuberculosis” OR “TB” OR “Mycobacterium tuberculosis”) AND (“drug resistance” OR “antimicrobial resistance” OR “AMR”) AND (“machine learning” OR “deep learning” OR “neural network”) AND (“prediction” OR “detection”) AND (“whole genome sequencing” OR “WGS” OR “SNPs” OR “genomics”). We screened the search results and performed a comprehensive literature review to identify suitable methods (see Supplementary File D1). Articles were included if they met the following criteria: (i) reported original research (excluding reviews), (ii) provided publicly available code, and (iii) presented no prohibitive implementation challenges. We identified 8 studies yielding 20 distinct models. Of these, seven were developed specifically for *M. tb*, while one was originally designed for *Escherichia coli* (*E. coli*) but adapted for this study. The methods were categorized into two major groups:

- **ML-based methods:** These include **Linear Models** (Logistic Regression [LR] with L1 and L2 regularization [LR L1 and LR L2], and LR with one-hot encoded [OHE] inputs [LR OHE]); **Tree-based methods** (Decision Tree, Random Forest [RF], RF with OHE, RF with phylogeny-related parallelism score (PRPS)-based inputs, Treesist, and XGBoost); **Support Vector Machines** (SVM with linear and RBF kernels and SVM with label-encoded [LE] and PRPS inputs); **Bayesian methods** (Bernoulli Naive Bayes); and **Shallow Artificial Neural Networks** (ANN) [9, 13–15].
- **DL-based methods:** These comprise Convolutional Neural Networks (CNN) with label-encoded inputs (CNN LE), DeepAMR, Wide and Deep Neural Networks (WDNN), Deep Neural Networks (Deep NN), and Single-Drug CNN (SDCNN) [7, 8, 16, 17].

Although our primary focus is on ML and DL methods, we also included the catalogue-based method TBProfiler [18] to serve as an established non-ML baseline.

#### 2.1.2 Feature Representation and Evaluation Pipeline

Most ML and ensemble-based approaches encode genomic variation — including single-nucleotide polymorphisms (SNPs) and insertions/deletions (INDELs) — as a binary feature matrix. In contrast, certain DL-based approaches directly model sequence data from resistance-associated genes. To account for this methodological heterogeneity, we evaluated the selected methods using three distinct input feature sets: (i) whole-genome variants, (ii) coding-region variants, and (iii) targeted variants or sequences from a subset of resistance-associated genes.

The evaluation pipeline begins with the preprocessing of WHO dataset samples (see Section 4.1), followed by the training of drug-specific models across the three feature sets. As illustrated in Figure 1, we employed different evaluation strategies to systematically assess performance. Table 1 summarizes the sample counts and feature dimensionality (SNPs and INDELs) available for each drug. Note that specific architecture required tailored inputs: PRPS-based models use a feature subset selected based on PRPS scores, LE/OHE-based methods use only SNP variants, and SDCNN uses drug-specific known resistance-associated gene sequences.

**Table 1.**
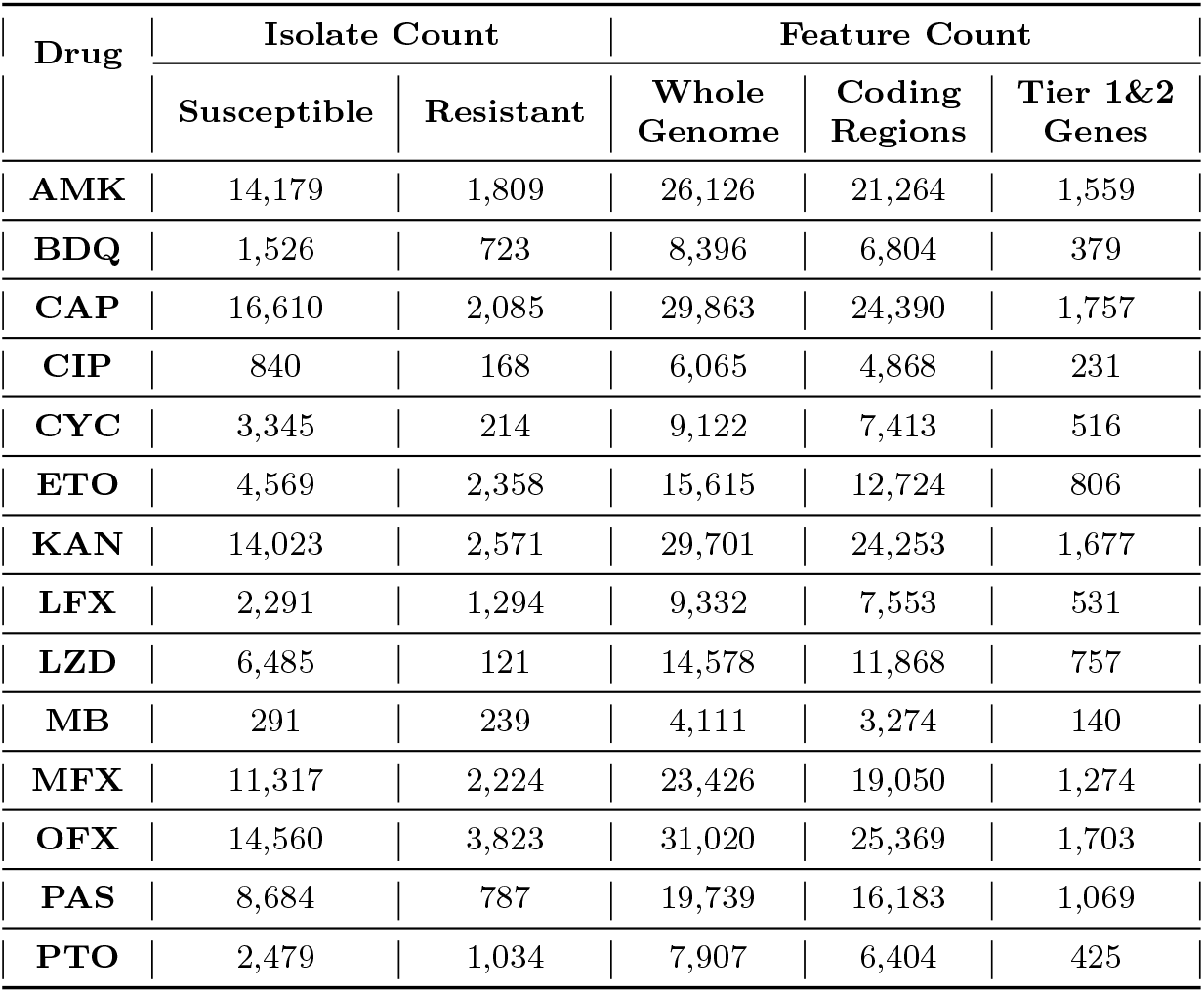
Distribution of sample sizes and feature dimensionality for the 14 second-line drugs: The table presents the counts of phenotypically susceptible and resistant isolates used for training and evaluation. It also details the number of input features (variants) extracted for each of the three feature sets: the entire genome, coding regions, and WHO Tier 1 and Tier 2 resistance-associated genes.

| Drug | Isolate Count |  | Feature Count |  |  |
| --- | --- | --- | --- | --- | --- |
|  | Susceptible | Resistant | Whole Genome | Coding Regions | Tier 1&2 Genes |
| AMK | 14,179 | 1,809 | 26,126 | 21,264 | 1,559 |
| BDQ | 1,526 | 723 | 8,396 | 6,804 | 379 |
| CAP | 16,610 | 2,085 | 29,863 | 24,390 | 1,757 |
| CIP | 840 | 168 | 6,065 | 4,868 | 231 |
| CYC | 3,345 | 214 | 9,122 | 7,413 | 516 |
| ETO | 4,569 | 2,358 | 15,615 | 12,724 | 806 |
| KAN | 14,023 | 2,571 | 29,701 | 24,253 | 1,677 |
| LFX | 2,291 | 1,294 | 9,332 | 7,553 | 531 |
| LZD | 6,485 | 121 | 14,578 | 11,868 | 757 |
| MB | 291 | 239 | 4,111 | 3,274 | 140 |
| MFX | 11,317 | 2,224 | 23,426 | 19,050 | 1,274 |
| OFX | 14,560 | 3,823 | 31,020 | 25,369 | 1,703 |
| PAS | 8,684 | 787 | 19,739 | 16,183 | 1,069 |
| PTO | 2,479 | 1,034 | 7,907 | 6,404 | 425 |

### 2.2 Performance of simple models versus complex DL methods

We compared the performance of models across three feature sets to identify approaches that demonstrated consistent robustness across varying input representations (see Section 4.1). Notably, the median feature dimensionality of the Tier 1 & 2 gene set is approximately 19-fold lower than that of the whole-genome set and 15-fold lower than that of the coding-region set. Despite this substantial reduction in dimensionality, models using the smaller feature sets perform comparably well to those trained on all variants (see Figure 2).

**Fig. 2.**
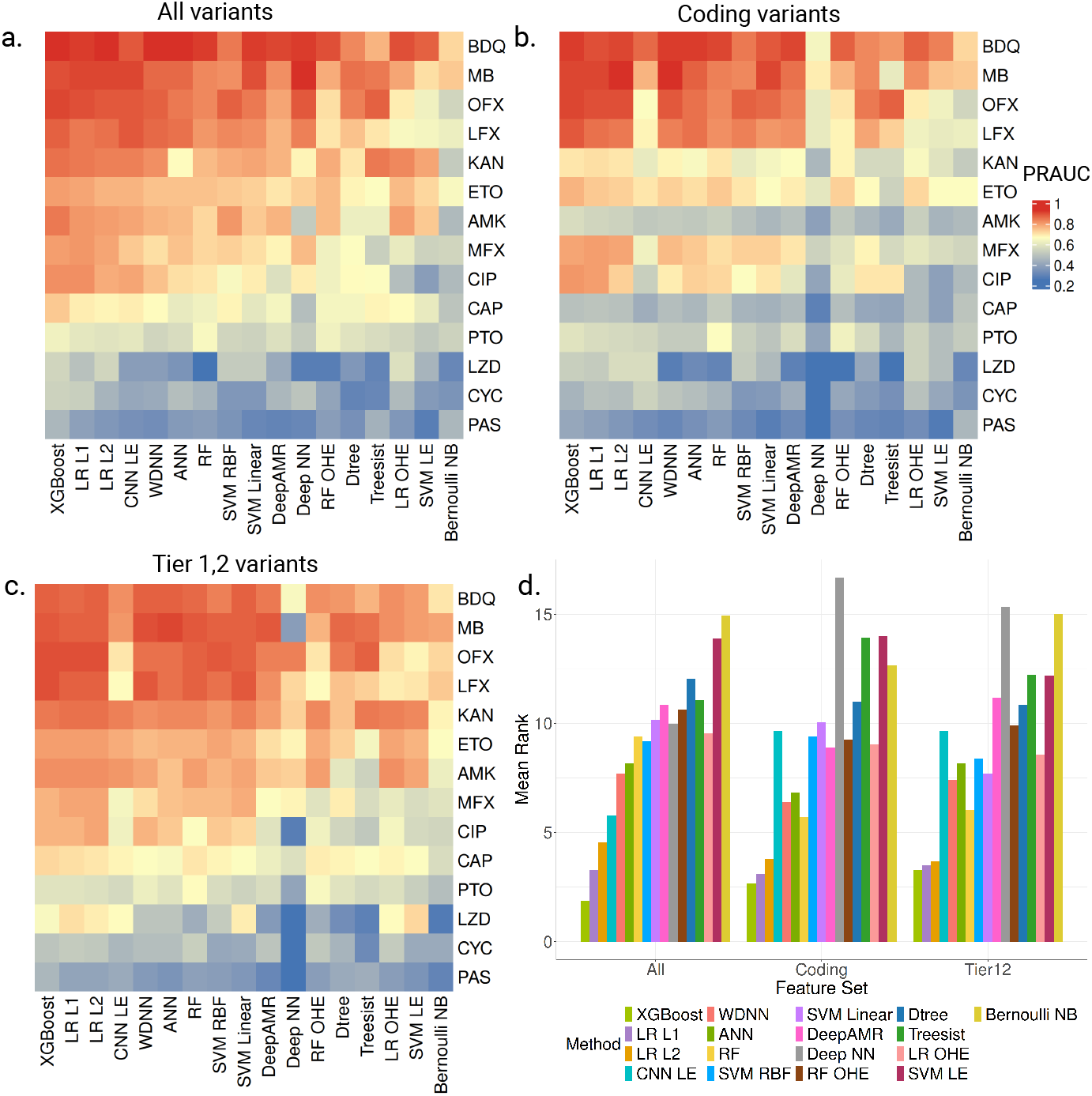
Comparison of PRAUC scores across methods and feature sets on held-out test data: a) Heatmap of PRAUC scores computed using genome-wide variants, and the methods and drugs are clustered using hierarchical clustering. b) Heatmap of PRAUC scores computed using variants from coding regions. c) Heatmap of PRAUC scores computed using variants from Tier 1&2 genes. The ordering of drugs and methods in panels b and c follows the ordering as in panel a. d) Bar chart showing overall method consistency across all drugs; lower summed ranks indicate best and consistent performance across most drugs.

#### 2.2.1 Performance on Whole-Genome Features

We first evaluated the PRAUC scores on the held-out test set for models trained on whole-genome features. For drugs such as AMK, BDQ, CAP, ETO, KAN, LFX, MB, and OFX, most models (excluding the Deep NN and Bernoulli NB) achieved high PRAUC values ranging from 60% to 95% (Figure 2a). Specifically, BDQ achieved a median PRAUC of 91%, while LFX, MB, and OFX showed median values between 80% and 90%. In contrast, models for MFX and PTO demonstrated moderate performance, with PRAUC values ranging from 50% to 81%. For drugs with severe class imbalance (resistant isolates < 10% of training data) — specifically, CYC, LZD, and PAS — no model exceeded a PRAUC of 60%, suggesting that data scarcity significantly hindered predictive performance.

#### 2.2.2 Impact of Feature Selection: Coding and Targeted Genes

When restricting inputs to coding region variants, performance dipped slightly. Median PRAUC values did not exceed 90% for any drug, although select models maintained high scores for BDQ, MB, and OFX. Drugs such as CIP, LFX, and MFX exhibited moderate performance, with a median PRAUC of 81% for LFX and 65–70% for CIP and MFX. For AMK, CAP, CYC, ETO, KAN, LZD, and PTO, median PRAUC values dropped to 40–70%. This reduction in performance relative to the whole-genome models may be attributed to the exclusion of resistant-conferring variants in non-coding regions, such as promoters [19].

Conversely, models trained on Tier 1 & 2 gene variants achieved PRAUC scores comparable to the whole-genome models, highlighting the efficacy of biologically curated feature sets. Interestingly, for LZD, the SVM (with LE input features) model achieved a PRAUC of 72% using Tier 1 & 2 variants — significantly outperforming all whole-genome models, none of which exceeded 60%. However, such improvements were not observed for the CYC and PAS drugs. Notably, the performance of the Bernoulli NB model also improved substantially when using Tier 1 & 2 variants compared to the all-variants case, likely due to a higher signal-to-noise ratio in the targeted gene set. Overall, these results indicate that simple ML models trained on well-curated features often outperform complex architectures, likely due to the specific structure of bacterial genomic data.

#### 2.2.3 Model Ranking and Selection

To identify the most robust models, we ranked models by PRAUC score for each drug and computed the mean rank across all feature sets (Figure 2d) XGBoost is the top-performing model across all feature sets, followed by LR L1 and LR L2. These results further underscore the ability of simple models to achieve higher precision–recall performance in predicting resistance to second-line drugs using binary variant data. Among the deep learning models, WDNN performs best, followed by CNN LE. Based on overall performance, XGBoost and LR-L1 emerged as the top-performing models within the ML category, while WDNN and CNN LE were the leading models within the DL category; these models were therefore selected for subsequent comparisons.

To quantify the impact of feature sets on these top models, we computed Δ_*gain*_, defined as the paired difference in PRAUC scores between models trained on each feature set. The whole-genome model showed only a marginal improvement over Tier 1 & 2 models (mean Δ_*gain*_ = 0.003 ±0.078). However, both the whole-genome and Tier 1 & 2 models demonstrated a clear advantage over the coding-region models (mean Δ_*gain*_ of 0.069 ±0.109 and 0.066 ±0.101, respectively). Given the comparable performance of the whole-genome models, we selected the top-performing whole-genome model for subsequent analyses.

*Note:* PRPS-based methods (SVC PRPS and RF PRPS) and SDCNN were excluded from the primary ranking analysis due to methodological constraints. PRPS-based models were evaluated for only eight drugs due to the computational challenges in generating PRPS trees for datasets exceeding 7,000 samples. SDCNN was excluded because its preprocessing pipeline removes samples with identical gene sequences, rendering the test sets inconsistent with the standard benchmark. However, PRAUC scores for these methods are available in Supplementary File D2; notably, their performance was generally comparable to the top-performing models across most drugs.

### 2.3 Performance profiling of top models

We next examined the performance characteristics of the top-performing ML and DL models to identify factors driving their predictive success. Comparing their PRAUC values against a random classifier — defined as the proportion of resistant isolates for each drug (Figure 3a) — reveals that both ML and DL models substantially out-perform random expectations. As the prevalence of resistant isolates increases, model performance also increases steadily up to approximately 0.20. Beyond this prevalence threshold, performance plateaus, suggesting that further increases in the proportion of resistant samples yield limited gains once this baseline is met.

**Fig. 3.**
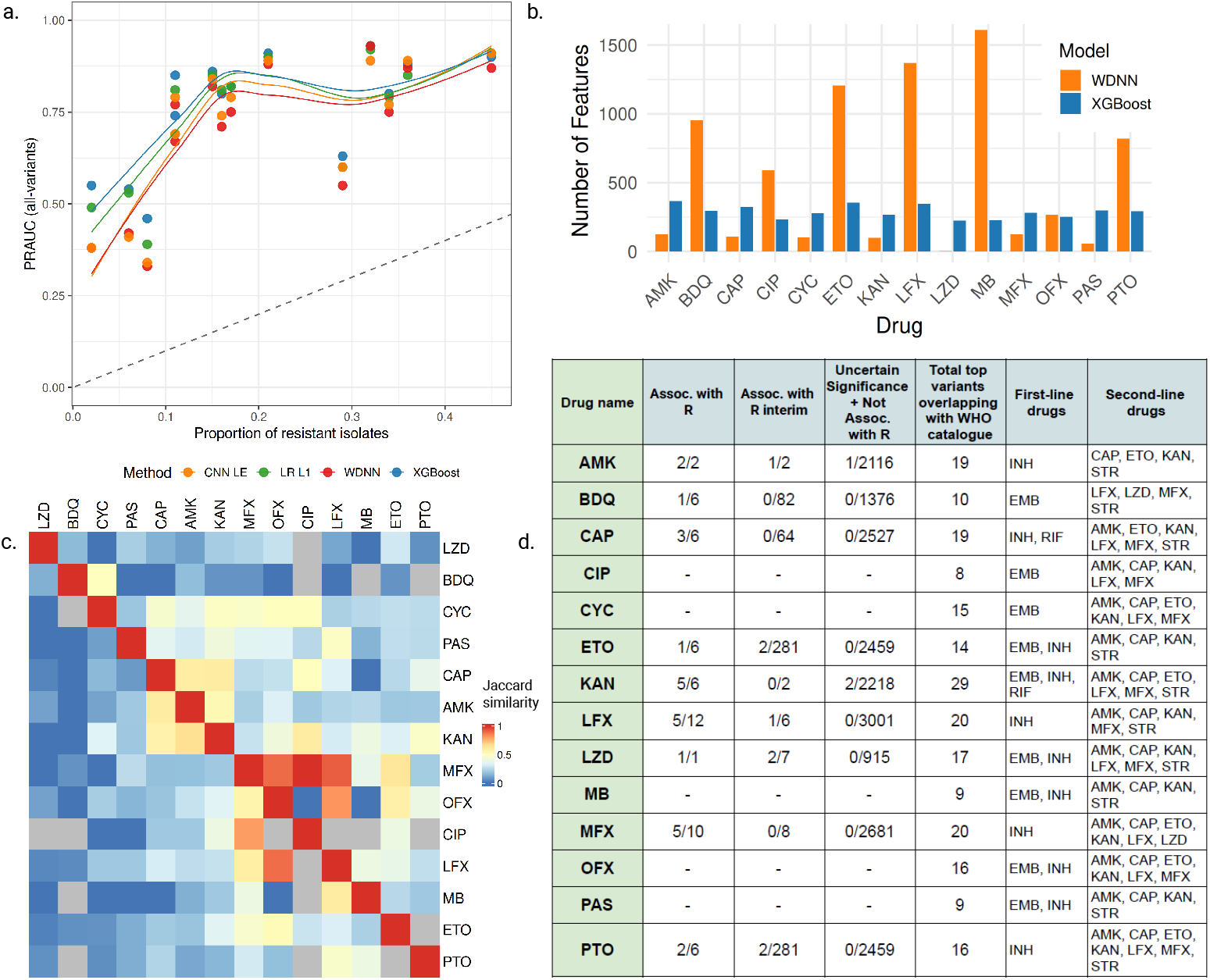
Performance and interpretability profiling of top models: a) Comparison of PRAUC scores of the top-performing models against the PRAUC of a random classifier. For each drug, the random classifier’s PRAUC corresponds to the proportion of resistant isolates observed for that drug. Number of top significant features identified by SHAP for XGBoost and WDNN. c) Heatmap showing the Jaccard similarity of true resistance labels between each drug pair (lower triangle) and the Jaccard similarity of predicted labels from XGBoost (upper triangle), computed using isolates from the held-out test set. d) Overlap between the top variants identified by XGBoost for each drug and those reported in the WHO 2023 catalogue. In entries formatted as a/b, a represents the number of top SHAP-identified variants, while b corresponds to the total number of variants reported in WHO catalogue for that category (can be ‘associated with resistance’, ‘associated with resistance interim’, ‘uncertain significance’, ‘not associated with resistance’ and ‘not associated with resistance interim’).

#### 2.3.1 Feature utilization and data sparsity

Given that the tree-based XGBoost model consistently outperformed the DL models, we further investigated differences in feature utilization between XGBoost and WDNN. We compared the top variants identified by SHapley Additive exPlanations (SHAP) at a cutoff >0.001 for both models under the whole-genome setting (Figure 3b). WDNN identified a larger number of significant features than XGBoost for most drugs. However, this trend was reversed for AMK, CAP, CYC, KAN, LZD, and PAS. Notably, for all these exceptions, the proportion of resistant isolates was ≤0.11. This suggests that the prediction task is inherently constrained by the scarcity of resistant samples, which likely limits the ability of complex models like WDNN to learn feature representations. In addition to data sparsity, cross-resistance mechanisms introduce further complexity, challenging the accurate identification of drug-specific resistance–associated features [20].

#### 2.3.2 Cross-resistance and co-occurrence patterns

Cross-resistance, a phenomenon where resistance to multiple drugs arises from shared biological mechanisms or overlapping causal variants, often results in highly correlated features within ML models. This correlation can cause models to attribute importance to spurious variants that reflect co-occurrence rather than true causality — an issue particularly relevant in TB, where cross-resistance between first-line and second-line drugs is frequent. [21]. To characterize the extent of this phenomenon in our dataset, we computed the Jaccard similarity for each pair of drugs using both the true phenotypic labels (resistant versus susceptible) and the labels predicted by the XGBoost model (see Figure 3c). The lower triangular matrix, representing the true labels, indicates that cross-resistance is generally low across most drug pairs. A similar pattern is observed in the upper triangular matrix, which represents the predicted labels. However, a few drugs — such as CYC, PAS, and MFX — exhibit slightly higher Jaccard similarity values in the predicted labels, suggesting that the model may be capturing underlying co-occurrence patterns or shared resistance mechanisms for these drugs. Given that such co-occurrence can complicate predictions, we next examined the interpretability of our models to assess their ability to correctly identify known resistance-associated mutations.

#### 2.3.3 Biological validation against the WHO catalogue

To validate the biological relevance of the learned features, we compared the top 50 variants identified by the XGBoost (whole-genome model) against the WHO mutation catalogue (see Supplementary File D2 for the top 50 variants). This comparison assessed whether the model prioritized known resistance-associated variants, including those linked to other first-line and second-line drugs. As shown in Figure 3d, the model successfully captured at least 50% of the high-confidence resistance-associated variants for most drugs, with the notable exception of BDQ and ETO. Crucially, the model also assigned high importance to variants known to confer resistance to other drugs. These findings confirm that cross-resistance can drive ML models to identify co-occurring variants as significant predictors, even when those variants are not mechanistically causal for the specific drug being analyzed.

#### 2.3.4 Ensembling and lineage-specific error analysis

To explore potential performance gains beyond individual models, we combined the predictions of the top-performing models, hypothesizing that different models might capture complementary resistance patterns. However, ensembling yielded no substantial improvement in performance (Supplementary Table S1). Finally, we analyzed the lineage distribution of samples misclassified by XGBoost and WDNN to detect potential lineage-specific biases. Lineages 2 and 4 were more predominant in the overall dataset, and the distribution of misclassified samples mirrored this prevalence. This suggests that classification errors are not driven by lineage-specific biases, but rather reflect the underlying distribution of lineages within the population (see Supplementary Figure S1).

### 2.4 Generalization performance

While the internal evaluation on the WHO dataset demonstrated strong performance across most drugs, translating this success to independent external validation datasets remains a challenge. We evaluated the top-selected models (trained on the WHO whole-genome variant set) on an external-validation dataset (see Section 4.1 for dataset details). Figure 4a, reveals a substantial drop in predictive power with PRAUC scores failing to exceed 75% for any drug.

**Fig. 4.**
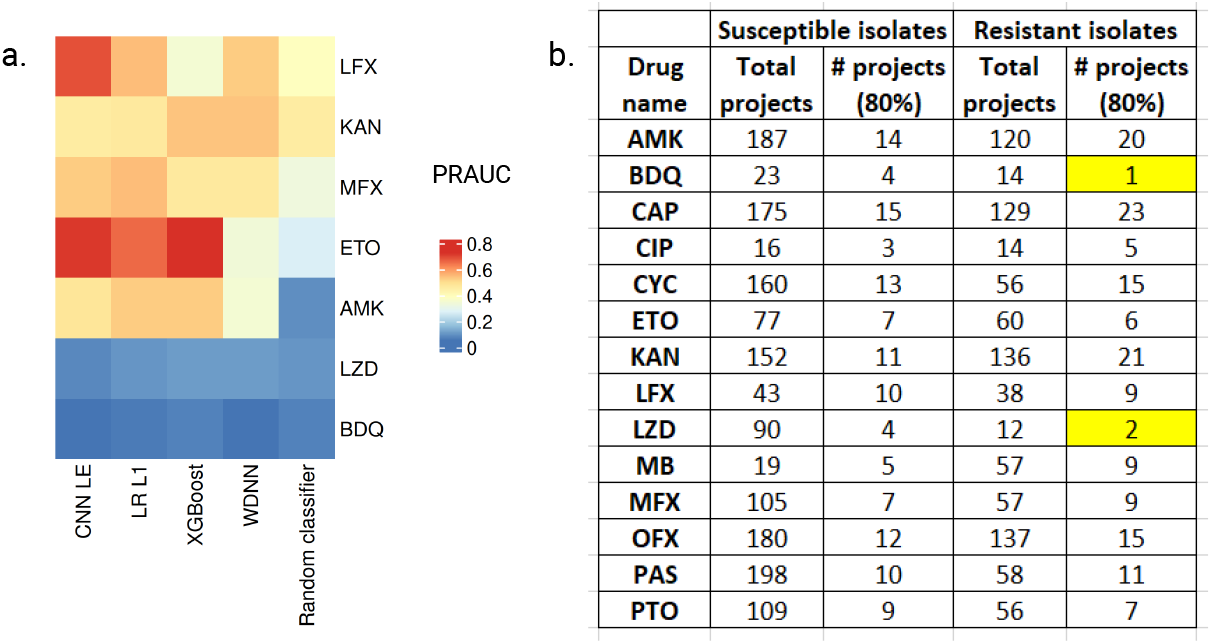
Performance comparison on the independent dataset and related analyses: a) Heatmap of PRAUC scores for the top-performing models trained on the WHO dataset (all-variants setup) and evaluated on the independent dataset. The corresponding values are provided in Supplementary Table S2. b) Table summarizing the distribution of project sources for samples corresponding to each drug in the WHO dataset. Approximately 80% of BDQ and LZD samples originated from a limited number of project sources (≤2), which are highlighted in yellow.

#### 2.4.1 Drug-Specific Performance

Among the top models, CNN LE achieved the highest PRAUC for LFX (69%), while XGBoost performed best for ETO (73%). The LR L1 model attained the top score only for MFX (55%). Notably, for BDQ and LZD, all models failed to outperform the random baseline (prevalence). Despite the fact that AMK — which has a similarly low resistance prevalence (<10% isolates) — maintained moderate predictive performance.

#### 2.4.2 Impact of Class Balancing

Based on these observations, hypothesizing that class imbalance might be driving this poor generalization, we retrained the top-performing XGBoost model using downsampling to balance susceptible and resistant isolates. While this strategy significantly improved performance on the internal held-out test set, it yielded no significant gains on the independent external validation dataset (Figure 5), suggesting class imbalance is not the primary driver of the generalization gap.

**Fig. 5.**
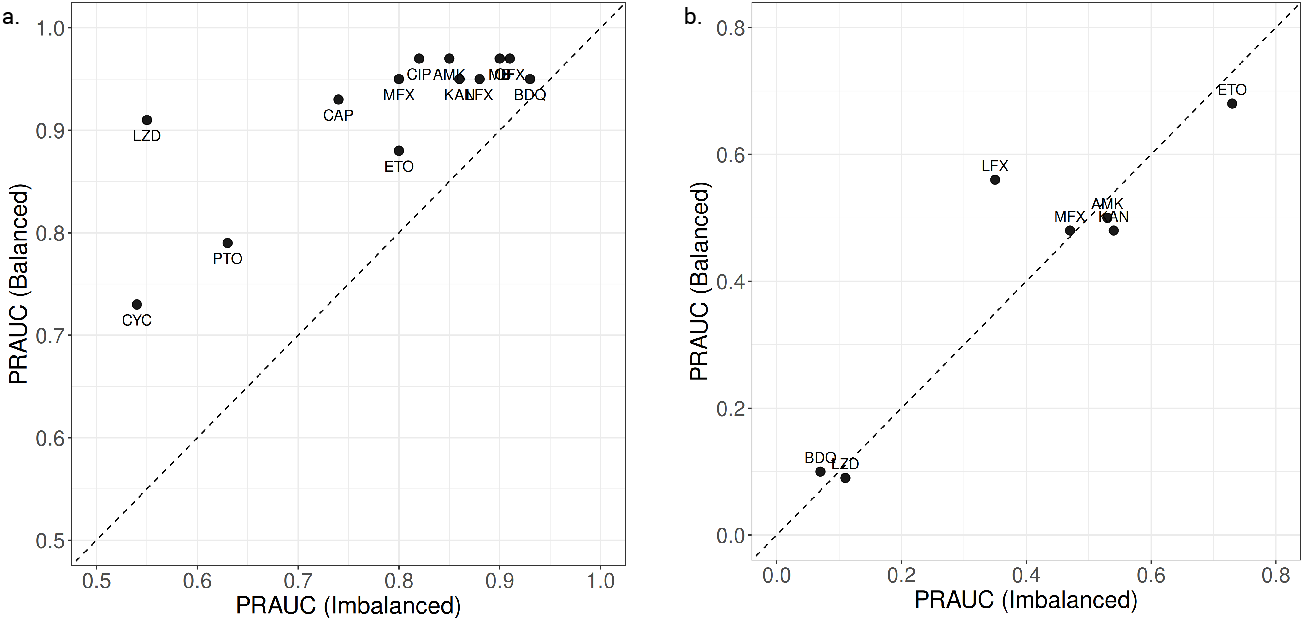
Performance comparison using balanced dataset: a) Scatter plot showing PRAUC scores of the XGBoost model (all-variants setup) evaluated on the held-out test set. b) Scatter plot showing PRAUC scores of the XGBoost model trained on balanced data (all-variants setup) and evaluated on the independent dataset. Here, imbalanced scores correspond to evaluations based on models trained using the original training dataset obtained from the WHO dataset split (see Section 4.2 for details).

#### 2.4.3 Dataset Bias and Confounding Factors

To investigate potential sources of this discrepancy, we analyzed the metadata of the WHO dataset, which is curated from multiple global studies (Figure 4b). This analysis revealed that approximately 80% of the samples for BDQ and LZD originate from one or two projects. This heavy skew suggests that the models likely learned study-specific or geographically specific features rather than globally relevant causal determinants [22]. These findings highlight the critical need for geographically diverse training data to ensure that computational models are robust to region-specific genomic variations before clinical deployment.

### 2.5 Comparison with catalogue-based TBProfiler

Catalogue-based methods, such as TBProfiler, rely on curated libraries of known resistance-associated variants to infer drug susceptibility. While conceptually simple, these approaches are robust, as the catalogue variants are validated through rigorous statistical association testing. To benchmark ML and DL models against this clinical standard, we evaluated TBProfiler on both the WHO held-out test set and the independent external dataset. Figure 6 compares the F1-scores of the top-performing models against TBProfiler (detailed results provided in Supplementary Files D3 and D4). In the held-out test set, TBProfiler generally performed comparably to, or slightly better than, the top ML/DL models. However, learned models demonstrated a performance advantage for specific drugs, notably LFX, MFX, and PTO. On the independent external dataset, TBProfiler exhibited poor performance for BDQ (10%), KAN (19%), and LZD (0%). While the WDNN model achieved a higher F1-score for KAN (54%) compared to TBProfiler, it still failed to outperform the random baseline (defined here as a classifier predicting all isolates as resistant). This highlights that while ML and DL methods can occasionally surpass catalogue-based tools for specific drugs, a significant gap remains in developing learnable models that consistently outperform expert-curated catalogues across the full spectrum of second-line drugs.

**Fig. 6.**
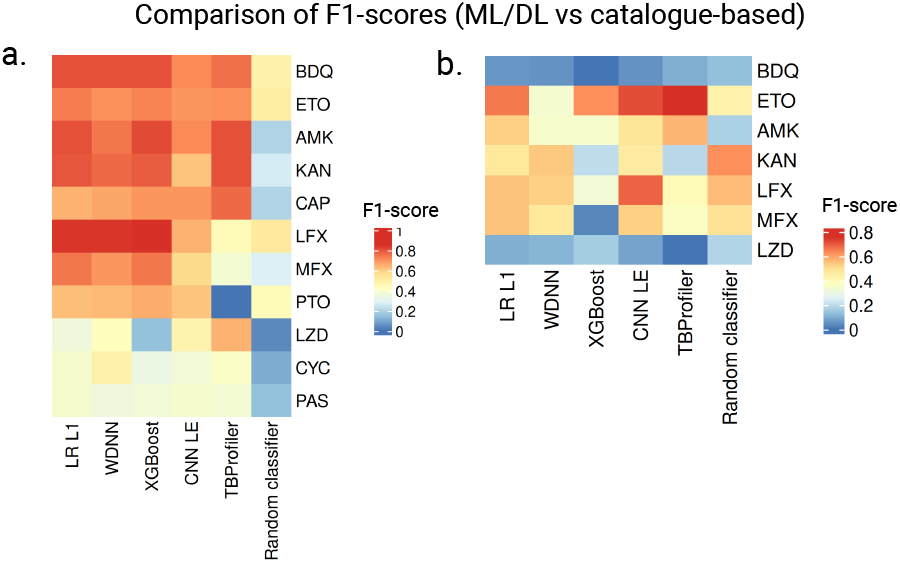
F1-scores of top models and the catalogue-based TBProfiler: a) Heatmap showing the F1-scores of the top models, TBProfiler, and random classifier on the WHO held-out test dataset. Note that the drugs CIP, MB, and OFX were excluded from the comparison, as TBProfiler did not provide predictions for these drugs in any of the samples. b) Heatmap showing the F1-scores of the top models alongside TBProfiler and random classifier on the independent dataset. The numerical values corresponding to both plots are presented in Supplementary Tables S3 and S4. Note that the random classifier corresponds to the all-positives classifier.

## 3 Discussion

We systematically surveyed the literature and comprehensively benchmarked 20 ML and DL methods for predicting TB drug resistance across 14 second-line drugs. The performance of all methods was evaluated on a large WHO dataset using multiple feature sets, and the top-performing models were further validated on an independent external validation dataset. Overall, our methods demonstrate that relatively simple ML models achieve robust performance, often rivaling or exceeding complex deep learning architectures even with reduced feature representations. Among the traditional ML approaches, XGBoost showed consistent superiority across all drugs, while WDNN emerged as the strongest DL model. However, despite comparable performance on the internal held-out test set, DL-based methods failed to demonstrate improved generalizability on the independent external dataset. Furthermore, our findings highlight the continued relevance of catalogue-based approaches, such as TBProfiler, which achieved predictive performance comparable to learnable methods, reinforcing the value of expert-curated knowledge in clinical diagnostics.

### 3.1 Model Complexity and Data Imbalance

Our analysis revealed that shallow models, such as XGBoost and Logistic Regression (LR), performed more robustly than complex DL models. This suggests that the current prediction task is driven primarily by the presence of dominant, high-effect resistance variants rather than the complex, non-linear multi-variant interactions that DL models are designed to capture. Consequently, these simpler models are well-suited for deployment in low-resource clinical settings where computational efficiency and interpretability are paramount.

### 3.2 The Impact of Sampling Bias and Generalization Gaps

A critical finding of this study is the disconnect between internal validation success and external generalization failure. For instance, moderate predictive performance (PRAUC <70%) was observed for drugs such as CYC, LZD, and PAS, likely due to severe class imbalance (<10% resistant samples). In contrast, BDQ, despite having a higher proportion of resistant samples (≈32%) and achieving the highest PRAUC on the held-out test dataset, showed poor generalization on the external validation dataset. This discrepancy suggests potential overfitting to dataset-specific patterns rather than robust resistance determinants. Indeed, our metadata analysis revealed that training data for BDQ and LZD originated largely from a limited number of project sources (≤ 2), suggesting the influence of sampling bias rather than intrinsic biological complexity. These results indicate that current large-scale datasets, while voluminous, may lack the population-level heterogeneity required to train globally generalizable models for newer second-line drugs. This underscores an urgent need for coordinated global initiatives to expand WGS cohorts, ensuring they capture diverse geographic and genetic backgrounds.

### 3.3 Feature Representation and Biological Insights

Interestingly, utilizing complex feature encoding schemes (e.g., One-Hot Encoding in RF/LR or Label Encoding in CNN/SVM) yielded no significant performance improvements over simple binary variant representations. Furthermore, models trained on curated, gene-specific feature sets performed comparably to those using genome-wide variants. This implies that for current sample sizes, the “signal” for resistance is concentrated in known loci, and adding genome-wide noise does not enhance predictive power.

Our analysis of feature importance also highlighted the challenge of cross-resistance. The overlap of top-ranked features across different drugs suggests that models may be learning shared markers of multi-drug resistance (MDR) or linear rather than drug-specific causal mechanisms. Lineage analysis confirmed that our samples were predominantly composed of Lineages 2 and 4; however, error analysis suggested that population structure was not the primary driver of misclassification. A key open question remains whether these shared features represent true pleiotropic mechanisms or merely confounding co-occurrence, warranting further investigation into integrative models that explicitly disentangle cross-resistance. Collectively, these findings suggest that relying solely on WGS variants may face an asymptote in performance, and integrating other omics data (e.g., transcriptomics) may be necessary to capture complex resistance phenotypes [23].

### 3.4 Limitations and Future Directions

While this study provides a comprehensive benchmark, several limitations remain. First, we did not explicitly quantify the impact of technical biases arising from different sequencing platforms or variant-calling pipelines, which may influence model generalizability. Future work should incorporate strategies to normalize these technical variations. Second, this study focused exclusively on binary classification (Resistant/Susceptible). Future benchmarking efforts should extend to quantitative prediction of minimum inhibitory concentration (MIC) values, which provide a more clinically refined measure of resistance. Finally, the inclusion of non-coding regulatory elements could further improve predictive resolution. Importantly, the generic computational framework established in this study is readily extensible, facilitating the standardized evaluation of emerging methods and datasets as they become available.

## 4 Methods

### 4.1 Datasets and preprocessing procedure

The genotypic and phenotypic data from the 2nd edition of the WHO mutation catalogue were used to train the predictive models [24]. The dataset comprised of 50,801 samples containing drug susceptibility information for 26 antibiotics (see Supplementary File D5 for biosample identifiers and related metadata). For samples with multiple sequencing runs, paired-end whole-genome sequencing (WGS) reads were prioritize over single-end runs.

We used a standardized in-house pipeline to preprocess each WGS sample and generate corresponding VCF files. Quality control was performed using fastp (v0.23.4) [25] to remove reads shorter than 50 base pairs, duplicates or those with an average quality score below 20. Filtered reads were aligned against the H37Rv reference genome (NCBI NC000962.3) using *BWA* (v0.7.18) [26]. Alignment duplicates were subsequently removed from the resulting BAM files using *Picard* (v2.20.4) [27]. Variant calling was executed with *freebayes* (v1.0.2) using default settings [28]. To ensure high-confidence calls for downstream analysis, variants were filtered using *bcftools* (v1.22) to retain only those with a depth (DP) ≥10 and an allele frequency (AF) ≥0.75 [29]. Here, AF represents the fraction of sequencing reads supporting the alternate allele at a specific variant position.

Variants were annotated using *SnpEff* (v5.2) [30] with only the top-ranked annotation retained for each variant. Synonymous variants and those located within PE/PPE gene families, including their Polymorphic GC-Rich Sequences (PGRS) subfamilies, were excluded from further analysis. Samples with an alignment mapping percentage below 90% or those derived from amplicon sequencing were also excluded, resulting in 49,266 high-quality samples (Supplementary File D6 lists the corresponding biosample identifiers).

From this filtered set, we focused on 14 second-line drugs that each possessed at least 100 resistant isolates. We subsequently selected 25,204 samples resistant to at least one of the 14 drugs for downstream analyses. For each drug, VCF files were merged using *bcftools* to generate a joint genotype, which was normalized and converted into a binary matrix (‘0’: absence of variant; ‘1:’ presence). This matrix was used to derive three distinct feature sets:

- **Genome-wide variant set:** Includes all variants observed across the entire genome.
- **Coding-region-specific variant set:** Includes all variants located within coding regions.
- **Resistance-gene-specific variant set:** Comprises variants located within the 73 Tier 1 and Tier 2 resistance genes utilized by `TBProfiler` (v4.4.0) [18].

All feature sets were further filtered to include only variants present in at least 10 samples. Note that the SVM-LE, LR-OHE, CNN-LE, and RF-OHE models utilized label or one-hot encoding for SNPs rather than binary indicators (see Section 4.3). Similarly, the SDCNN model used gene sequences from known resistance-associated genes as input instead of individual variants.

To evaluate generalization performance, we utilized an independent external validation dataset from China, comprising 9,071 samples [31]. We selected only isolates resistant to at least one of the 7 available second-line drugs (AMK, BDQ, ETO, KAN, LFX, LZD, or MFX), resulting in a validation subset of 1,199 samples (see Supplementary File D7 for details). The distribution of the number of resistant samples chosen for analysis from this dataset is given in Table 2.

**Table 2.** Sample size distribution for the independent validation dataset: The table details the number of phenotypically susceptible and resistant isolates available for each drug in the external validation cohort, used to assess generalization.

| Drug | Susceptible | Resistant |
| --- | --- | --- |
| AMK | 1,088 | 111 |
| BDQ | 262 | 19 |
| ETO | 874 | 325 |
| KAN | 650 | 546 |
| LFX | 735 | 464 |
| LZD | 253 | 27 |
| MFX | 805 | 394 |

### 4.2 Training and testing

For each drug, we developed drug-specific models using an 80/20 stratified split (80% training+validation, 20% held-out test set) to ensure that the original class imbalance was preserved in both the training and testing sets. Models were trained on the WHO dataset using either predefined static parameters or optimal hyperparameters identified through systematic tuning (refer to Section 4.3 and Supplementary File D6 for specific parameter settings).

Once the model parameters were fixed, we determined the optimal decision threshold for class assignment using four-fold cross-validation on the training set. This threshold was selected by maximizing the average Youden’s J statistic across folds to balance sensitivity and specificity.

The final models were then trained on the complete training set (the 80% split) using the selected threshold and subsequently evaluated on the held-out test set. Predictive performance was primarily assessed using the area under the precision-recall curve (PRAUC) and the F1-score, computed for both training and test predictions (see Section 4.4 for details on evaluation metrics used).

### 4.3 Background on benchmarked methods

This section details the implementation of the 20 ML and DL architectures bench-marked in this study. Comprehensive parameter settings for all models are provided in Supplementary File D8. Due to implementation constraints and preliminary testing that showed marginal gains from extensive search, hyperparameter tuning was restricted to DeepAMR and PRPS-based methods. For all other architectures, we adopted the default parameter configurations used by the original authors.

#### 4.3.1 ML-based models

The ML-based model are categorized into binary-input-based and encoding-based models, as described below:

##### Binary-input-based models

These models use a binary feature matrix as input, with entries of 0 or 1 indicating the absence or presence of a variant during training. The variant set includes both SNPs and INDELs.

- **LR L1, LR L2, SVM Linear, SVM RBF, RF, Bernoulli NB:** These models were selected based on a study that employed traditional methods for resistance prediction and originally used a dataset comprising 1,839 samples [13] for the analysis. All models were implemented using the corresponding classifiers from the *scikit-learn* package.
- **Dtree and Treesist:** Treesist was proposed as an extension of the decision tree (Dtree) framework, integrating prior biological knowledge through feature prioritization, biologically informed tree pruning, and gene-level constraints [9]. In the original study, Treesist was compared with the catalogue-based TBProfiler and a standard decision tree (Dtree) using 32,689 samples. In our benchmarking analysis, we incorporated prior information indicating whether variants originated from coding or non-coding regions when applying Treesist and also included Dtree as one of the models for comparison. Dtree was implemented using the corresponding classifier from *scikit-learn* package, whereas Treesist employs a modified splitting criterion to determine the best split. In the all-variants setting, the coding or non-coding status of each variant was used as prior information.
- **XGBoost and ANN:** These models were based on the recent study that applied them to predict resistance to 5 first-line and 8 second-line TB drugs using WGS data from 13,947 samples [14]. The ANN employed is a simple architecture with two hidden layers using Rectified Linear Unit (ReLU) and Sigmoid activation functions, respectively. The model was trained using parameters suggested by the authors, except that the batch size and number of training epochs were set to 32 and 50, respectively.
- **SVC PRPS and RF PRPS:** The study introduces a novel phylogeny-related PRPS, a metric that quantifies the extent to which a feature’s distribution reflects underlying population structure [15]. By filtering features strongly associated with phylogeny, the integration of PRPS with SVM and RF models reduces the feature space while improving predictive performance.

#### Encoding-based models

Instead of using binary features, the study by Ren et al. proposed three encoding strategies, namely, OHE, LE, and Frequency Chaos Game Representation (FCGR) for training the ML models LR, SVM, and RF [16]. These encodings incorporate either the reference or alternate nucleotides at each variant position and therefore consider only SNPs. OHE represents each variant as a five-dimensional binary vector corresponding to (N, A, G, C, T), while label encoding maps nucleotides to integer values. FCGR transforms genome sequences into two-dimensional fractal representations using the chaos game representation. In the original paper, the ML models were applied to encodings constructed from WGS data of *E. coli*. The authors have also used a CNN model for their analysis, which is discussed in the next section under deep learning models. For benchmarking purposes, a single encoding was selected for each method using a ranking criterion based on four-fold cross-validation on the training dataset. For each method–drug pair, the encoding that achieved the highest PRAUC on the validation data was counted as a win, and the encoding with the maximum number of wins was chosen as the optimal representation for that model. Based on this analysis, LE was selected for the SVM and CNN models, while OHE was chosen for the RF and LR models.

#### 4.3.2 DL-based models

Static parameters were used to train all deep learning models, except DeepAMR. Additional method-specific details are provided below.

##### Deep NN and WDNN

In [7], the authors used a WDNN model that combines a wide logistic regression component with a deep neural network comprising three hidden layers of 256 ReLU units with dropout, batch normalization, and L2 regularization, and a Deep NN with the same architecture but without the wide component. The dataset used for analysis consists of 3,601 samples.

##### DeepAMR

This deep denoising autoencoder model was used to predict resistance to first-line drugs and was trained on a dataset comprising 8,388 isolates [17]. The architecture is a symmetric autoencoder with successive fully connected layers that compress the input data into a low-dimensional latent representation of size 20, with Gaussian dropout applied for regularization. We also used the authors’ implementation to select the optimal learning rate using a cyclic learning rate algorithm. In addition, we determined the best learning rate across four folds using 80% of the training data and selected the final learning rate based on the average PRAUC performance across the folds.

##### CNN

These are widely used for classifying two-dimensional images and have also found applications in the analysis of biological data. Ren et al. [16] have applied a CNN consisting of eleven hidden layers, including convolutional, batch normalization, pooling, flattening, fully connected, and dropout layers to predict drug resistance using encoded genomic data (as discussed in Section 4.3.1).

##### SDCNN

A deep convolutional neural network (CNN) was trained on genome sequences from drug-specific, known resistance-associated coding regions to predict antimicrobial resistance [8]. In this benchmarking study, only single-drug models were used for the analysis.

To identify the most informative features used by the models for prediction, we employed the SHapley Additive exPlanations (SHAP) framework in Python. For the top-performing models, XGBoost and WDNN, we utilized the model-specific SHAP explainers, TreeExplainer and GradientExplainer, respectively, to compute normalized importance scores for individual variants. Only features with scores greater than 0.001 were retained for downstream analysis.

### 4.4 Evaluation measures

We used the following quantitative measures to optimize model parameters and evaluate predictive performance:

#### Youden’s J statistic

This metric was used to identify the optimal decision threshold by maximizing the vertical distance from the diagonal line on the Receiver Operating Characteristic (ROC) curve. It is defined as:

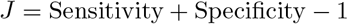

where 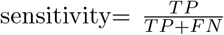 and 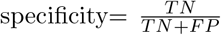. Here, *TP, TN, FP*, and *FN* represent the counts of true positives, true negatives, false positives, and false negatives, respectively. This statistic was computed during the four-fold cross-validation phase to calibrate the decision threshold.

##### Precision–Recall Area Under the Curve (PRAUC)

PRAUC was used to evaluate predictive performance, particularly under class imbalance. The precision–recall curve illustrates the trade-off between precision and recall across varying decision thresholds, and PRAUC provides a single summary measure corresponding to the area under this curve. Recall is equivalent to sensitivity, and precision is defined as:

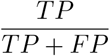

Higher PRAUC values indicate better discriminative performance, especially for the minority class.

#### F1-score

The F1-score summarizes precision and recall by computing their harmonic mean:

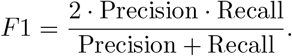

This metric provides a balanced measure of performance when both false positives and false negatives are important.

#### Jaccard similarity

Jaccard similarity quantifies the overlap between two sets of elements *A* and *B* and is defined as:

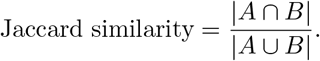

It ranges from 0 (no overlap) to 1 (complete overlap).

## Data and code availability

All source code for data preprocessing, the generic framework for model training and evaluation, and the associated analysis scripts are available on GitHub: https://github.com/BIRDSgroup/TB-Bench. Raw WGS sequencing data for the samples corresponding to the WHO mutation catalogue and the external validation dataset are publicly available in the NCBI Sequence Read Archive (SRA) and the Genome Sequence Archive (GSA), respectively. The accession identifiers for samples from the WGS and independent datasets are provided in Supplementary Data Files D6 and D7, respectively. All additional supplementary files and source data have been deposited in https://drive.google.com/drive/folders/17DBf0hSLbbl0xsRwQb_7DQhSCP1qok6r?usp=drive_link.

## Acknowledgements

We thank members of our Bioinformatics and IntegRative Data Science (BIRDS) group for their valuable input during the presentations of this work. This work is supported by the Women Leading IIT Madras (WLI) grant (SB24250033CSIITM008892) awarded to BVP, and Prime Minister’s Research Fellowship (PMRF) grant (SB22230881CSPMRF003119) awarded to SJ.

## Notes

https://github.com/BIRDSgroup/TB-Bench

### Competing Interest Statement

The authors have declared no competing interest.

