## Supplementary Information (Supplementary Tables/Figures/Files) for "TB-Bench: A Systematic Benchmark of Machine Learning and Deep Learning Methods for Second-Line TB Drug Resistance Prediction"

### 1 Supplementary Tables

**Table S1**: **Comparison of top-performing models and their ensemble predictions on the held-out test dataset:** The PRAUC scores of the four top-performing models and their ensemble predictions on the WHO held-out test dataset are given. For ensemble predictions, resistance was assigned when two or more individual classifiers predicted the resistant class.

| Drug | XGBoost | LR L1 | WDNN | CNN LE | Ensemble |
|---|---|---|---|---|---|
| AMK | 0.85 | 0.81 | 0.77 | 0.79 | 0.75 |
| BDQ | 0.93 | 0.92 | 0.93 | 0.89 | 0.88 |
| CAP | 0.74 | 0.69 | 0.67 | 0.69 | 0.66 |
| CIP | 0.82 | 0.82 | 0.75 | 0.79 | 0.75 |
| CYC | 0.54 | 0.53 | 0.42 | 0.41 | 0.37 |
| ETO | 0.80 | 0.79 | 0.75 | 0.77 | 0.72 |
| KAN | 0.86 | 0.85 | 0.82 | 0.84 | 0.79 |
| LFX | 0.88 | 0.85 | 0.87 | 0.89 | 0.85 |
| LZD | 0.55 | 0.49 | 0.38 | 0.38 | 0.31 |
| MB | 0.90 | 0.91 | 0.87 | 0.91 | 0.81 |
| MFX | 0.80 | 0.81 | 0.71 | 0.74 | 0.75 |
| OFX | 0.91 | 0.90 | 0.88 | 0.89 | 0.88 |
| PAS | 0.46 | 0.39 | 0.33 | 0.34 | 0.39 |
| PTO | 0.63 | 0.60 | 0.55 | 0.60 | 0.63 |

**Table S2**: **Generalization performance:** The PRAUC scores for the top-performing models and the random classifier on the independent dataset are presented.

| **Models** | **XGBoost** | **LR L1** | **CNN LE** | **WDNN** | **Random classifier** |
|---|---|---|---|---|---|
| **AMK** | 0.53 | 0.53 | 0.48 | 0.35 | 0.09 |
| **BDQ** | 0.07 | 0.06 | 0.05 | 0.05 | 0.07 |
| **ETO** | 0.73 | 0.66 | 0.72 | 0.34 | 0.27 |
| **KAN** | 0.54 | 0.47 | 0.46 | 0.54 | 0.46 |
| **LFX** | 0.35 | 0.55 | 0.69 | 0.53 | 0.39 |
| **LZD** | 0.11 | 0.10 | 0.08 | 0.11 | 0.1 |
| **MFX** | 0.47 | 0.55 | 0.53 | 0.47 | 0.33 |

**Table S3**: **Comparison of top models with TBProfiler on the held-out test dataset:** Drug-specific F1-scores of the top-performing models and TBProfiler evaluated on the WHO held-out test dataset.

| Drug Name | XGBoost | LR L1 | CNN LE | WDNN | TBProfiler | Random classifier |
|---|---|---|---|---|---|---|
| **AMK** | 0.82 | 0.81 | 0.72 | 0.75 | 0.81 | 0.2 |
| **BDQ** | 0.81 | 0.81 | 0.72 | 0.81 | 0.76 | 0.49 |
| **CAP** | 0.7 | 0.65 | 0.7 | 0.67 | 0.77 | 0.2 |
| **CYC** | 0.34 | 0.39 | 0.37 | 0.49 | 0.41 | 0.11 |
| **ETO** | 0.73 | 0.74 | 0.7 | 0.71 | 0.71 | 0.51 |
| **KAN** | 0.79 | 0.8 | 0.62 | 0.77 | 0.81 | 0.27 |
| **LFX** | 0.86 | 0.85 | 0.65 | 0.85 | 0.45 | 0.53 |
| **LZD** | 0.15 | 0.36 | 0.48 | 0.43 | 0.65 | 0.04 |
| **MFX** | 0.75 | 0.75 | 0.58 | 0.7 | 0.38 | 0.28 |
| **PAS** | 0.37 | 0.39 | 0.39 | 0.36 | 0.38 | 0.15 |
| **PTO** | 0.66 | 0.63 | 0.62 | 0.64 | 0 | 0.45 |

**Table S4**: **Comparison of top models with TBProfiler on the independent dataset:** Drug-specific F1-scores of the top-performing models and TBProfiler evaluated on the independent dataset.

| Drug Name | XGBoost | LR L1 | CNN LE | WDNN | TBProfiler | Random classifier |
|---|---|---|---|---|---|---|
| **AMK** | 0.35 | 0.53 | 0.48 | 0.35 | 0.57 | 0.17 |
| **BDQ** | 0.00 | 0.06 | 0.05 | 0.05 | 0.1 | 0.13 |
| **ETO** | 0.63 | 0.66 | 0.72 | 0.34 | 0.76 | 0.43 |
| **KAN** | 0.20 | 0.47 | 0.46 | 0.54 | 0.19 | 0.63 |
| **LFX** | 0.33 | 0.55 | 0.69 | 0.53 | 0.4 | 0.56 |
| **LZD** | 0.16 | 0.10 | 0.08 | 0.11 | 0 | 0.18 |
| **MFX** | 0.03 | 0.55 | 0.53 | 0.47 | 0.37 | 0.49 |

#### 2 Supplementary Figures

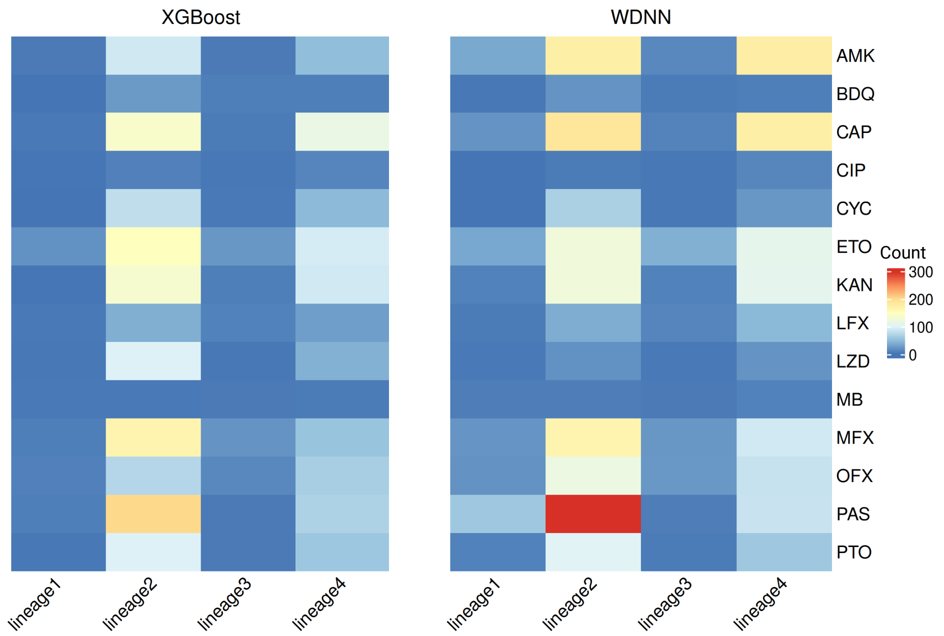


**Fig. S1**: **Lineage-specific misclassification counts across drugs by top-performing models:** Heatmap shows the number of samples misclassified by both XGBoost and WDNN within each lineage for each drug.

### 3 Supplementary Files

All additional supplementary files and source data have been deposited in https://drive.google.com/drive/folders/17DBf0hSLbbl0xsRwQb_7DQhSCP1qok6r?usp=drive_link.

**Supp File D1:** This file provides detailed information on the articles considered for the systematic review. The "inclusion status" field specifies whether each article was included in or excluded from the final evaluation.

**Supp File D2:** This file contains the PRAUC and F1 scores of the models evaluated on the WHO held-out test dataset across 14 drugs using three feature sets (all variants, coding variants, and Tier 1&2 variants). Results are organized in separate tabs, and an additional tab provides the top 50 variants identified through SHAP analysis.

**Supp File D3:** TBProfiler-predicted drug resistance profiles for 49,264 samples from the WHO mutation catalogue dataset, including drug-specific resistance calls used for F1-score calculation in our analysis. A sample was considered resistant to a given drug if it contained at least one associated resistance-conferring variant.

**Supp File D4:** TBProfiler-predicted drug resistance profiles for samples in the independent dataset, including drug-specific resistance calls used for F1-score calculation in our analysis. A sample was considered resistant to a given drug if it contained at least one associated resistance-conferring variant.

**Supp File D5:** This file contains the metadata corresponding to the raw data of the WHO samples, including biosample identifiers. The number of rows exceeds the number of samples (50,801), as some samples have duplicates due to multiple sequencing.

**Supp File D6:** This file contains the biosample identifiers of the 49,266 samples retained after filtering. Among these, 25,204 samples are resistant to at least one second-line drug.

**Supp File D7:** This file contains the accession numbers of samples downloaded from the GSA repository that are resistant to at least one second-line drug.

**Supp File D8:** This file provides the fixed parameter settings used for training the 20 models.
